# Towards next generation diagnostics for tuberculosis: identification of novel molecular targets by large-scale comparative genomics

**DOI:** 10.1101/569384

**Authors:** Galo A. Goig, Manuela Torres-Puente, Carla Mariner-Llicer, Luis M. Villamayor, Álvaro Chiner-Oms, Ana Gil-Brusola, Rafa Borrás, Iñaki Comas

## Abstract

Tuberculosis remains one of the main causes of death worldwide. The long and cumbersome process of culturing *Mycobacterium tuberculosis* complex (MTBC) bacteria has encouraged the development of specific molecular tools for detecting the pathogen. Most of these tools aim to become novel tuberculosis diagnostics, and big efforts and resources are invested in their development, looking for the endorsement of the main public health agencies. Surprisingly, no study had been conducted where the vast amount of genomic data available is used to identify the best MTBC diagnostic markers. In this work, we use large-scale comparative genomics to provide a catalog of 30 characterized loci that are unique to the MTBC. Some of these genes could be targeted to assess the physiological status of the bacilli. Remarkably, none of the conventional MTBC markers is in our catalog. In addition, we develop a qPCR assay to accurately quantify MTBC DNA in clinical samples.

## Background

Tuberculosis (TB) is the most lethal infectious disease caused by a single agent, namely bacteria belonging to the *Mycobacterium tuberculosis* complex (MTBC)[1]. Whereas isolating the bacteria from clinical specimens is a time-consuming process that delays both clinical diagnosis and research workflows, rapid molecular tests have the potential to identify the pathogen DNA in a few hours [2,3]. This is the main reason why the development of new molecular tools for TB diagnosis is an active area of research, with many companies involved, looking for the endorsement of the World Health Organization (WHO) [4]. The most successful example has been the Xpert MTB/RIF test [5], which was endorsed by the WHO back in 2010 for TB diagnosis, and recommended as the first-line diagnostic in 2017[6]. Achieving a high sensitivity and specificity is pivotal for the development and improvement of molecular tests to ensure an accurate diagnosis. To this end, most tests incorporate specific markers for the detection of MTBC bacteria. For instance, the new Xpert MTB/RIF Ultra assay, previously targeting the *rpoB* gene alone, has now incorporated the insertion sequences IS6110 and IS1081[7]. The insertion sequence IS6110 has been extensively used as a MTBC-specific marker since first described in 1990[8]. In addition, the IS6110 can be present in high copy numbers in some MTBC strains (from 0 to 27 copies)[9], causing the nucleic acid amplification tests (NAAT) targeting this sequence to achieve higher sensitivities for strains carrying several copies. However, the specificity of the IS6110 has been questioned since two decades ago[10–15] what, along with the fact that some strains lack this insertion sequence, can lead to an incorrect diagnosis[16,17].

Several other genes have been used as markers for the accurate identification of MTBC bacteria[18–21]. However, the accuracy of NAATs based on these markers rely on the specificity of the primers, since most of the targeted loci are claimed to be MTBC-specific, yet they were evaluated with limited genomic information on the diversity of NTM and MTBC bacteria.

Nowadays, the use of the publicly available *omic* data can help identifying species-specific genetic markers to develop accurate molecular tools. Analyzing *omic* data has been proven to be an effective strategy for the identification of specific markers in several organisms[22–26], and even some workflows have been published for the evaluation of genetic markers based on genomic data[27]. For instance, comparative genomics was used by Zozaya-Valdés *et al.* to assess the population structure of *Mycobacterium chimaera*, identifying six specific loci of these organisms that allowed them to develop a highly accurate qPCR assay.

Strikingly, the use of comparative genomics for the identification of MTBC-specific loci has been very limited. The few published studies focused on genetic regions acquired by horizontal gene transfer and used the limited datasets available at the time of publication, a decade ago[28–30]. By contrast, last years have witnessed a burst of available genomic sequences of a wide range of mycobacteria species and thousands of strains of the MTBC[31–33].

In this work, we perform a large-scale comparative genomic analysis to provide a reference list of 30 MTBC-specific loci that will be of great utility for the scientific community working on the development of new research and clinical tools for tuberculosis. Remarkably, we found that the main MTBC markers used up to date are also present in other organisms, mainly NTM. In our analysis, we assess the global diversity of each MTBC-specific gene among a comprehensive dataset of more than 4,700 MTBC strains, showing the value of using the genomic data at hand to identify the best targets for diagnostic assays. In addition, we develop a qPCR assay based on one of these markers capable of quantifying MTBC DNA in clinical samples.

## Methods

### In silico identification of MTBC-specific diagnostic gene markers

To identify MTBC-specific loci, we used blastn[34] to look for all the genes of the tuberculosis reference strain H37Rv (NC_000962.3) in the NCBI nucleotide non-redundant database (accessed October 2018) and a custom database comprising 4,277 NTM assemblies (Supplementary Methods 1). All the searches were performed specifying the algorithm blastn with a word size (or seed) of 7 bp. Then, we filtered the results with a set of stringent parameters to discard loci similar to any genomic region of any organism other than MTBC. We discarded all the genes that presented an alignment of more than 25% of its sequence (query coverage) with a similarity greater than 80%. If a gene was aligned in 60% of its sequence or longer it was discarded regardless of the similarity of the alignment. We only kept those genes that were present in all the MTBC bacteria.

Once potential MTBC-specific markers were identified, we decided to assess their genetic diversity. To do this, we analyzed the polymorphisms (single nucleotide polymorphisms (SNPs) and indels) observed at each position across a dataset comprising 4,766 genomes of MTBC strains[35]. Therefore, the number of SNPs of each gene was calculated as the sum of positions showing any nucleotide other than the reference. In the case of indels, we considered those positions showing an indel in at least 10 strains (0.2% of the database) to avoid the noise introduced by single-strain indels spanning large genic regions and possible false deletions arising as a result of sequencings with uneven genomic coverages. This allowed us to calculate different metrics for each gene such as the absolute number of polymorphisms, polymorphisms per base and, most importantly, the prevalence of each one.

Finally, we looked for available information of these genes in the bibliography, what allowed us to discard some candidates based on their genomic context and provide extended information about their physiology. We gathered transcriptomic and proteomic data derived from different published studies: transcriptomic data in response to overexpression of 206 transcription factors[36], different genotoxic stresses[37] and response to nitric oxide stress at different time-points[38], as well as proteomic data in response to nutrient starvation[39].

### Set-up of a MTBC-specific qPCR assay for DNA detection and quantification

We used the list of 30 MTBC-specific loci to set up a qPCR assay for the detection and quantification of MTBC DNA. To select the target for the assay, we took into consideration the number of polymorphisms per base, the absence of high-prevalent polymorphisms, the gene length and its genomic context. These criteria enabled an optimum design of primers, amplifying a universal and highly-specific region for the detection of MTBC. We designed the primers and probes for the assay using the web tool Primer-BLAST[40], checking that no unspecific amplicons were predicted. Finally, the qPCR assay consisted on the amplification of a 65 bp region within the Rv2341 gene using the following primers: Forward-GCCGCTCATGCTCCTTGGAT, Reverse-AGGTCGGTTCGCTGGTCTTG, Probe-TGAGTGCCTGCGGCCGCAGCGC.

To test the specificity of the assay we performed qPCR experiments with DNA from all MTBC lineages (except lineage 7 due to unavailability), human DNA, a mock sample with mixed DNA from 20 different bacterial species (ATCC^®^ MSA-1002^™^ and 17 different species of NTM (Supplementary Methods 2).

The reaction efficiency was calculated using serial dilutions of pure H37Rv DNA as template (0.5 ng/ul to 0.5*10^-5^ ng/ul). In addition, we evaluated the performance of the assay detecting and quantifying MTBC DNA in a test set of clinical samples. We used extracted DNA from 12 homogenized sputum samples from culture-positive TB patients, two of them with negative smear microscopy. We also used a DNA extraction from a non-TB patient sputum to spike in known concentrations of pure H37Rv DNA (0.5 ng/ul to 0.5*10^-5^ ng/ul), to calculate the reaction efficiency in clinical samples.

All the qPCR reactions were carried out using hydrolysis probes chemistry (FAM/BHQ) in a total volume of 20ul, containing 10ul of Kapa Probe Fast Master Mix 2X (Kapa Biosystems), 250mM of each primer, 350mM of probe and 2ul of sample. All were performed in a Roche Lightcycler 96 (Roche Diagnostics), with two replicates per sample and including reactions with no template as negative controls (NTC). When calculating reaction efficiencies, we used three replicates per point instead of two. The conditions for each assay comprised an initial denaturation step at 95°C for 3 minutes, followed by 55 amplification cycles as follows: 20 seconds at 60°C for annealing, 1 second at 72°C for extension, and 10 seconds at 95°C for denaturation. The results were analyzed with LightCycler 96 ® 1.1 software. Triplicates of each assay were carried out to check the reproducibility.

### Bacterial culture, clinical specimens and DNA extraction

All the DNA extractions were performed in our laboratory except for the commercial DNA mix of 20 bacterial species. Available cultures of different NTM species were subcultured in in 7H11 solid agar media and then the DNA extracted following the standard CTAB protocol[41] with an inactivation step of 1 hour at 80°C. DNA concentrations were measured with the Qubit fluorometer (dsDNA high-sensitivity kit) and samples with a concentration higher than 1ng/ul were normalized to 1ng/ul. In the case of the 13 sputum specimens, DNA extraction was performed as described by Votintseva *et al*[42]. All the samples were handled in a BSL-3 until DNA was extracted and purified.

### Ethics approval

The clinical specimens used in this study were collected as part of the surveillance program of communicable diseases by the General Directorate of Public Health of the Comunidad Valenciana and, as such, falls outside the mandate of the corresponding Ethics Committee for Biomedical Research. All personal information was anonymized and no data allowing individual identification was retained.

## Results

We identified 40 genes to be uniquely present in members of the MTBC according to our filtering parameters (Figure 1). After evaluating their genetic diversity across a database of more than 4,700 MTBC strains, we observed that the median number of SNPs per base was 0.07, with some of these genes showing either higher or lower diversities (up to 0.1 and 0.04 SNPs/base respectively), probably as a result of different selective pressures. Importantly, although most of the polymorphisms analyzed were strain-specific, we observed high prevalent polymorphisms as well (Figure 1, Supplementary File 1). For instance, Rv0610c showed a SNP present in 4182 strains and Rv2823c showed an insertion in 4,345 strains. Analysis of the phylogenetic distribution of these polymorphisms confirmed that they mapped to deep branches in the phylogeny. For example, the SNP in Rv0610c affected all modern lineages (L2, L3, L4).

**Figure 1.**
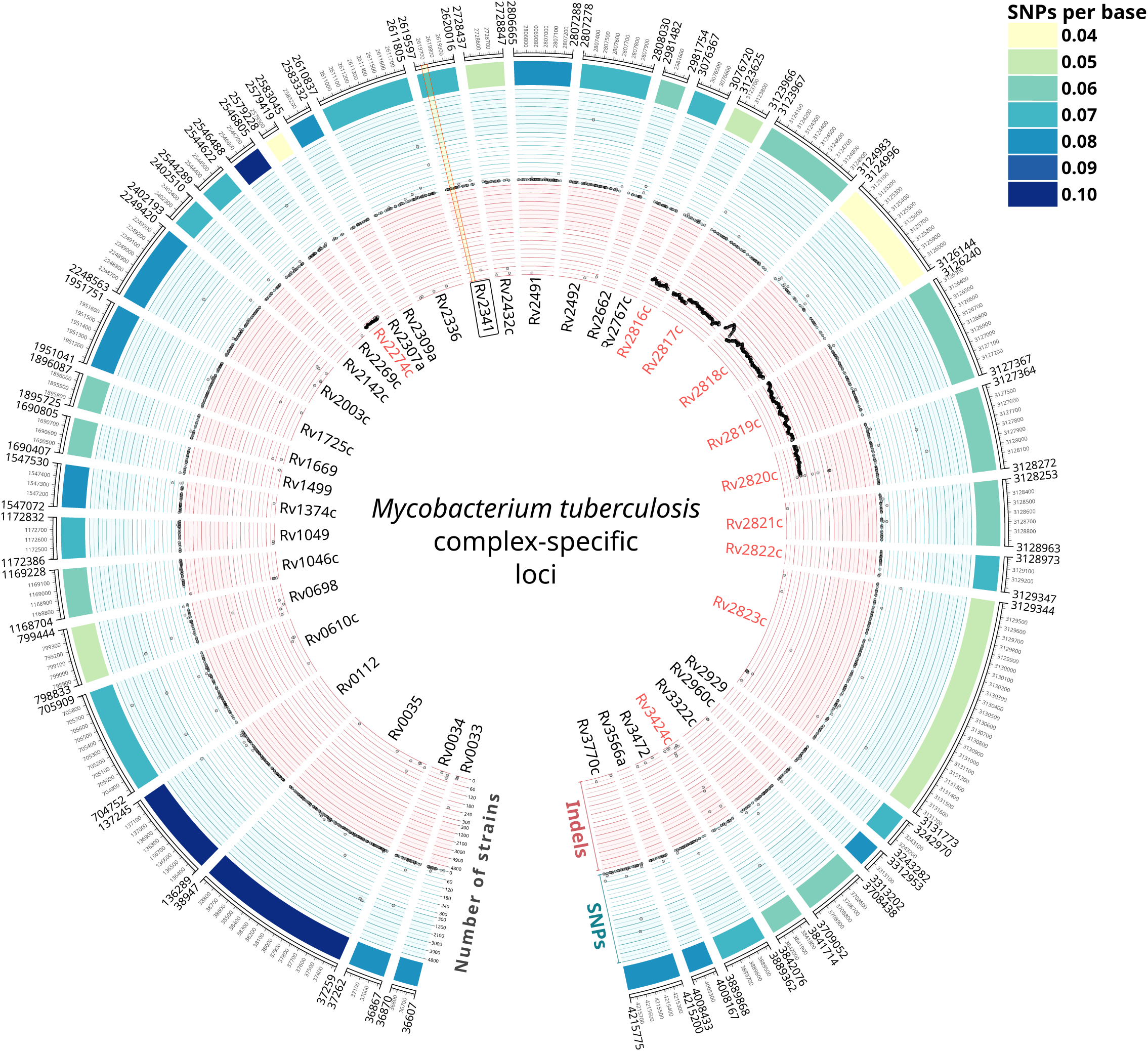
The 40 *Mycobacterium tuberculosis* complex (MTBC)-specific loci identified after an extensive search with blast in the NCBI non-redundant nucleotide database and a custom database of 4,277 Non-tuberculous mycobacteria (NTM). Gene names in red indicate loci that were discarded as diagnostic markers for being within regions of difference (Rv2274c within RD 182 and Rv2816c-2820c within RD 207), associated to CRISPR (Rv2816c-2823c) or duplicated in the genome (Rv3424c). Concentric circles represent genetic diversity metrics calculated by analyzing a dataset of 4,766 MTBC strains. Outer circle: heatmap representing the number of SNPs per base. Blue circle: prevalence of each SNP of each gene across the database of MTBC strains. Inner, read circle: prevalence of each indel of each gene across the database of MTBC strains. Note that both inner circles have two scales, one from 0 to 300 strains and other from 300 to 4,800 strains. The region of the Rv2341 gene amplified in our qPCR assay, avoiding prevalent polymorphisms, is indicated in light yellow. Note that regions of difference 182 and 207 are clearly detected in our analysis, indicated as contiguous deleted regions in a high number of strains.

Among these, 9 genes were discarded as potential diagnostic markers since they were included in regions of difference (RD) 182 (Rv2274c) and RD 207 (Rv2816c-Rv2820c) as described in Gagneux *et al.*[43] or were in variable genomic regions associated to CRISPR elements (Rv2816c-2823c)[44]. Another gene, Rv3424c was also discarded as we found it to be duplicated in a very labile genomic region, between the (putative) transposase of the insertion sequence IS1532 and PPE 59. Therefore, the curated list of MTBC-specific diagnostic markers finally consisted in 30 genes (Figure 1).

When looking at published transcriptomic and proteomic data (see Methods), we observed that Rv2003c, Rv2142c, and Rv3472 proteins are found in greater levels (6.19, 3.6 and 100-fold respectively) when the bacteria is subjected to starvation. Interestingly, Rv2003c is also observed to be overexpressed upon treatment with nitric oxide (Supplementary File 2).

Based on our large genomic analysis, we set up a qPCR assay targeting the Rv2341 gene. This gene, described as “probable conserved lipoprotein lppQ” in the Mycobrowser database[45], is situated in a stable genomic region, between the asparagine tRNA and the gene of the DNA primase, involved in the synthesizes of the okazaki fragments. Furthermore, we were able to design an optimized set of primers that avoid, at the same time, any region harboring prevalent polymorphisms (Figure 1).

When testing the qPCR assay with a panel of samples including different MTBC lineages, human, mock bacterial communities and different NTMs, the specificity of the assay was of 100%. The efficiency of the reaction was of 95% showing a limit of detection of 10fg (hypothetically corresponding to 2 genome equivalents). When using a standard curve of pure H37Rv DNA spiked in sputum samples, both the efficiency of the reaction (97%) and the limit of detection remained unaltered (Figure 2). When testing our qPCR assay with a panel of 12 TB sputum samples, we were able to detect and quantify MTBC DNA in all TB patient sputa, including 2 confirmed TB cases with a negative smear microscopy (Supplementary File 4).

**Figure 2.**
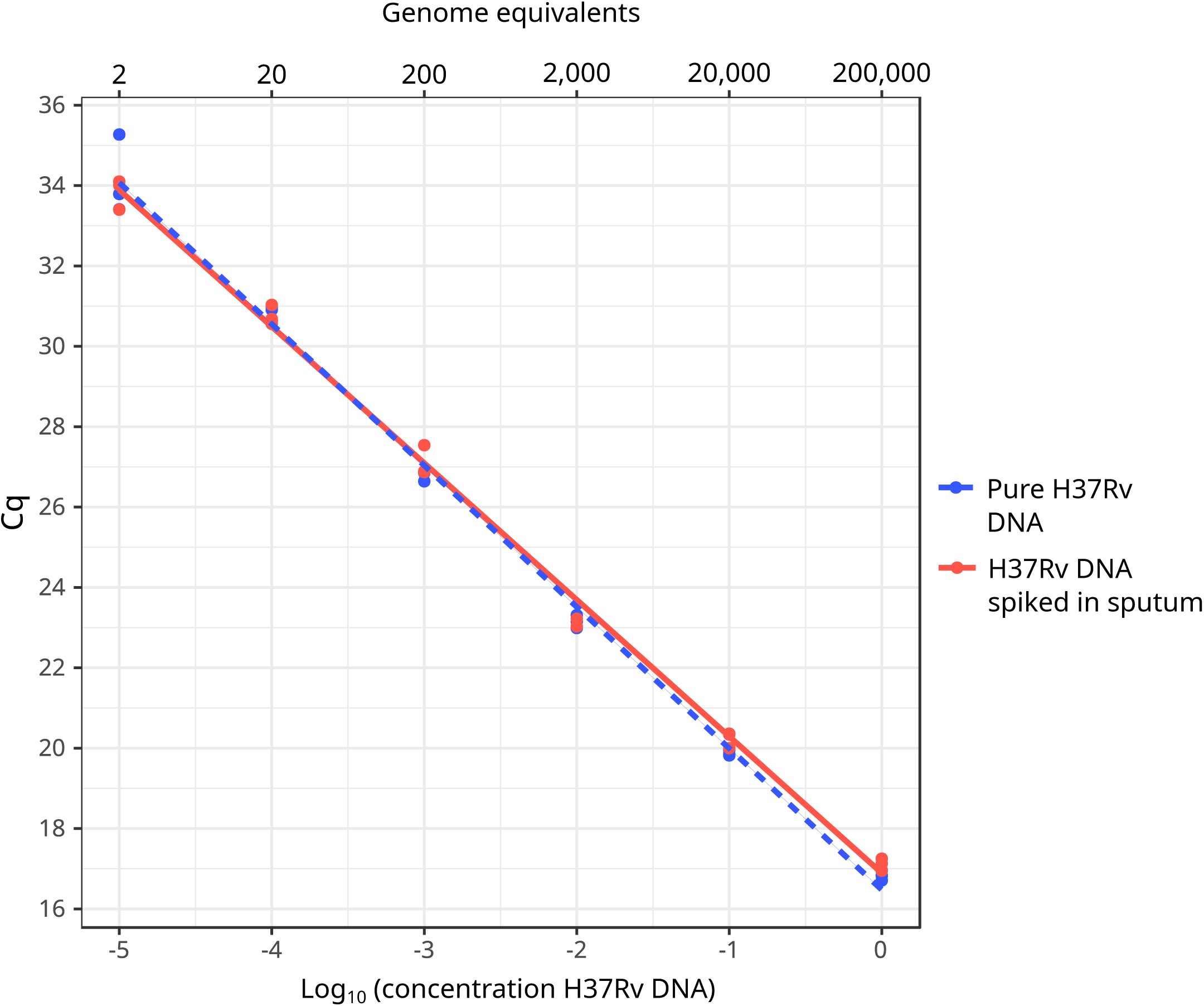
Standard curve for the qPCR assay targeting Rv2341 using known quantities of pure H37Rv DNA (in blue; efficiency=95%) and pure H37Rv DNA spiked in sputum samples (in red; efficiency=97%). In the upper x-axis is represented the hypothetical number of genome copies. All qPCR experiments were carried out in triplicates to check for reproducibility.

## Discussion

Identification of MTBC markers for the development of new diagnostic and research tools for tuberculosis has been an active area of research over the last decades, focusing on the direct or indirect detection of the tubercle bacilli. It is striking that for such a relevant disease, from both the epidemiological and economical point of view, for which tons of genomic data is already available, the identification of MTBC-specific genes had been relegated to the background. This has been probably motivated by the fact that current molecular tools have shown to perform well in most of situations. For instance, assays targeting the insertion sequence IS6110 ([46] or *rpoB*[47]. However, the available tools are not enough to stop the spread of the disease and for this reason many new generation diagnostics are still being developed with the aim to improve the accuracy of the existing ones and tackle their known flaws.

Our analysis provides invaluable information to develop such diagnostics, with a catalog of specific MTBC markers. Remarkably, some of the markers that we identify could be targeted to determine the physiological status of MTBC bacteria under certain conditions. For example, Rv2003c, overexpressed during starvation and upon treatment with nitric oxide[38,39], is also upregulated during dormancy[48]. Similarly, Rv1374c has been described to be a small RNA that is highly expressed during exponential growth[49], and hence could be used to evaluate the replicative state of the bacilli.

Strikingly, none of the markers considered to be MTBC-specific up to date are in our list of unique MTBC genes. For instance, when examining in which species the IS6110 can be found, we observed several non-MTBC organisms, including 14 NTMs, carrying at least one copy. The same is true for IS1081 and *mpt64*, present in 38 and 6 NTM respectively (Supplementary File 3). Similarly, the short-chain dehydrogenase/reductase gene (SDR) (Rv0303, region 365,234–366,142), which has been recently described as a *M. tuberculosis*-specific marker[28], is actually present in several NTM, as revealed by a blastn search in the non-redundant database of the NCBI web server (accessed January 2019), and in our database of NTM assemblies (Supplementary File 5). The fact that IS6110 is still one of the most used genetic targets for MTBC DNA detection (for example in the new Xpert Ultra MTB/RIF assay[7]), highlights the great utility, and the necessity, of translating the results of genomic analyses to the laboratory.

To illustrate the translational potential of our work, we set up an accurate qPCR assay capable of quantifying MTBC DNA with 100% specificity and a sensitivity up to 2 genome copies. Quantifying MTBC DNA from clinical samples is challenging due to the presence of PCR inhibitors along with great proportions of DNA from human and oropharyngeal microbiota. However, this capability is invaluable not only for diagnostic purposes, but also in the research context, for example when developing new protocols[42,50]. Remarkably, our assay, targeting a small region of the Rv2341 gene, showed an excellent performance in a test set of clinical specimens. However, we want to highlight that the list provided here comprehends 30 loci, from which many different molecular tools for tuberculosis could be developed.

Altogether, our analysis has a direct translational value, as it represents an important resource for research groups and companies involved in the development and improvement of novel TB diagnostics. For instance, the markers identified in this work could be used to improve existing tests such as the Xpert MTB/RIF assay, by including targets that we have demonstrated to be globally conserved and fully specific to the MTBC.

## Declarations

### Author Contributions

GAG and IC designed the study and analyzed the data. GAG and ACO analyzed the 4,766 MTBC strains dataset. GAG, MTP and CML performed the qPCR experiments. GAG and LMV cultured the non-tuberculous mycobacteria and performed the DNA extractions. AGB and RB did the microbiological identification of the isolates of non-tuberculous mycobacteria. RB provided the clinical specimens and clinical data. GAG and IC wrote the first draft of the manuscript. All authors contributed to the final version of the manuscript.

## References

1. World Health Organization. Global Tuberculosis Report 2017. 2017.

2. Eddabra R, Benhassou HA. Rapid molecular assays for detection of tuberculosis. Pneumonia. BioMed Central. 2018; 10(1):4.

3. Machado D, Couto I, Viveiros M. Advances in the molecular diagnosis of tuberculosis: From probes to genomes. Infect Genet Evol. 2018; Available from: http://dx.doi.org/10.1016/j.meegid.2018.11.021

4. Pai M, Nicol MP, Boehme CC. Tuberculosis Diagnostics: State of the Art and Future Directions. Microbiol Spectr. 2016; 4(5). Available from: http://dx.doi.org/10.1128/microbiolspec.TBTB2-0019-2016

5. Cirillo DM, Miotto P, Tortoli E. Evolution of Phenotypic and Molecular Drug Susceptibility Testing. Adv Exp Med Biol. 2017; 1019:221–246.

6. WHO meeting report of a technical expert consultation: non-inferiority analysis of Xpert MTF/RIF Ultra compared to Xpert MTB/RIF. Geneva: World Health Organization; 2017 (WHO/HTM/TB/2017.04); Available from: http://www.who.int/tb/publications/2017/XpertUltra/en/

7. Dorman SE, Schumacher SG, Alland D, et al. Xpert MTB/RIF Ultra for detection of *Mycobacterium tuberculosis* and rifampicin resistance: a prospective multicentre diagnostic accuracy study. Lancet Infect Dis. 2018; 18(1):76–84.

8. Thierry D, Brisson-Noël A, Vincent-Lévy-Frébault V, Nguyen S, Guesdon JL, Gicquel B. Characterization of a Mycobacterium tuberculosis insertion sequence, IS6110, and its application in diagnosis. J Clin Microbiol. 1990; 28(12):2668–2673.

9. Roychowdhury T, Mandal S, Bhattacharya A. Analysis of IS6110 insertion sites provide a glimpse into genome evolution of *Mycobacterium tuberculosis*. Sci Rep. 2015; 5:12567.

10. Kent L, McHugh TD, Billington O, Dale JW, Gillespie SH. Demonstration of homology between IS6110 of *Mycobacterium tuberculosis* and DNAs of other *Mycobacterium spp*. J Clin Microbiol. 1995; 33(9):2290–2293.

11. Liébana E, Aranaz A, Francis B, Cousins D. Assessment of genetic markers for species differentiation within the *Mycobacterium tuberculosis* complex. J Clin Microbiol. 1996; 34(4):933–938.

12. McHugh TD, Newport LE, Gillespie SH. IS 6110 homologs are present in multiple copies in mycobacteria other than tuberculosis-causing mycobacteria. J Clin Microbiol. 1997; 35(7):1769–1771.

13. Hellyer TJ, DesJardin LE, Beggs ML, et al. IS6110 homologs are present in multiple copies in mycobacteria other than tuberculosis-causing mycobacteria. J Clin Microbiol. 1998; 36(3):853–854.

14. Müller R, Roberts CA, Brown TA. Complications in the study of ancient tuberculosis: non-specificity of IS6110 PCRs. STAR: Science & Technology of Archaeological Research. 2015; 1(1):1–8.

15. Coros A, DeConno E, Derbyshire KM. IS 6110, a *Mycobacterium tuberculosis* complex-specific insertion sequence, is also present in the genome of *Mycobacterium smegmatis*, suggestive of lateral gene transfer among mycobacterial species. J Bacteriol. 2008; 190(9):3408–3410.

16. Steensels D, Fauville-Dufaux M, Boie J, De Beenhouwer H. Failure of PCR-Based IS6110 analysis to detect vertebral spondylodiscitis caused by *Mycobacterium bovis*. J Clin Microbiol. 2013; 51(1):366–368.

17. Huyen MNT, Tiemersma EW, Kremer K, et al. Characterisation of *Mycobacterium tuberculosis* isolates lacking IS6110 in Viet Nam. Int J Tuberc Lung Dis. 2013; 17(11):1479–1485.

18. Therese KL, Jayanthi U, Madhavan HN. Application of nested polymerase chain reaction (nPCR) using MPB 64 gene primers to detect *Mycobacterium tuberculosis* DNA in clinical specimens from extrapulmonary tuberculosis patients. Indian J Med Res. 2005; 122(2):165–170.

19. Chakravorty S, Sen MK, Tyagi JS. Diagnosis of extrapulmonary tuberculosis by smear, culture, and PCR using universal sample processing technology. J Clin Microbiol. 2005; 43(9):4357–4362.

20. Nimesh M, Joon D, Pathak AK, Saluja D. Comparative study of diagnostic accuracy of established PCR assays and in-house developed sdaA PCR method for detection of *Mycobacterium tuberculosis* in symptomatic patients with pulmonary tuberculosis. J Infect. 2013; 67(5):399–407.

21. Queipo-Ortuño MI, Colmenero JD, Bermudez P, Bravo MJ, Morata P. Rapid differential diagnosis between extrapulmonary tuberculosis and focal complications of brucellosis using a multiplex real-time PCR assay. PLoS One. 2009; 4(2):e4526.

22. Carmona SJ, Sartor PA, Leguizamón MS, Campetella OE, Agüero F. Diagnostic peptide discovery: prioritization of pathogen diagnostic markers using multiple features. PLoS One. 2012; 7(12):e50748.

23. Buchanan CJ, Webb AL, Mutschall SK, et al. A Genome-Wide Association Study to Identify Diagnostic Markers for Human Pathogenic *Campylobacter jejuni* Strains. Front Microbiol. 2017; 8:1224.

24. Carrera M, Böhme K, Gallardo JM, Barros-Velázquez J, Cañas B, Calo-Mata P. Characterization of Foodborne Strains of *Staphylococcus aureus* by Shotgun Proteomics: Functional Networks, Virulence Factors and Species-Specific Peptide Biomarkers. Front Microbiol. 2017; 8:2458.

25. Wang H, Drake SK, Yong C, et al. A Genoproteomic Approach to Detect Peptide Markers of Bacterial Respiratory Pathogens. Clin Chem. 2017; 63(8):1398–1408.

26. Koul S, Kumar P. A Unique Genome Wide Approach to Search Novel Markers for Rapid Identification of Bacterial Pathogens. J Mol Genet Med. 2015; 09(04).

27. Felten A, Guillier L, Radomski N, Mistou M-Y, Lailler R, Cadel-Six S. Genome Target Evaluator (GTEvaluator): A workflow exploiting genome dataset to measure the sensitivity and specificity of genetic markers. PLoS One. 2017; 12(7):e0182082.

28. Kakhki RK, Neshani A, Sankian M, Ghazvini K, Hooshyar A, Sayadi M. The short-chain dehydrogenases/reductases (SDR) gene: A new specific target for rapid detection of *Mycobacterium tuberculosis* complex by modified comparative genomic analysis. Infect Genet Evol. 2019; Available from: http://dx.doi.org/10.1016/j.meegid.2019.01.012

29. Becq J, Gutierrez MC, Rosas-Magallanes V, et al. Contribution of horizontally acquired genomic islands to the evolution of the tubercle bacilli. Mol Biol Evol. 2007; 24(8):1861–1871.

30. Veyrier F, Pletzer D, Turenne C, Behr MA. Phylogenetic detection of horizontal gene transfer during the step-wise genesis of *Mycobacterium tuberculosis*. BMC Evol Biol. 2009; 9:196.

31. Fedrizzi T, Meehan CJ, Grottola A, et al. Genomic characterization of Nontuberculous Mycobacteria. Sci Rep. Nature Publishing Group; 2017; 7:45258.

32. Coll F, Phelan J, Hill-Cawthorne GA, et al. Genome-wide analysis of multi- and extensively drug-resistant Mycobacterium tuberculosis. Nat Genet. Nature Publishing Group; 2018; 50(2):307.

33. CRyPTIC Consortium and the 100,000 Genomes Project, Allix-Béguec C, Arandjelovic I, et al. Prediction of Susceptibility to First-Line Tuberculosis Drugs by DNA Sequencing. N Engl J Med. 2018; 379(15):1403–1415.

34. Altschul SF, Gish W, Miller W, Myers EW, Lipman DJ. Basic local alignment search tool. J Mol Biol. 1990; 215(3):403–410.

35. Chiner-Oms Á, González-Candelas F, Comas I. Gene expression models based on a reference laboratory strain are poor predictors of *Mycobacterium tuberculosis* complex transcriptional diversity. Sci Rep. 2018; 8(1):3813.

36. Turkarslan S, Peterson EJR, Rustad TR, et al. A comprehensive map of genome-wide gene regulation in *Mycobacterium tuberculosis*. Sci Data. 2015; 2:150010.

37. Namouchi A, Gómez-Muñoz M, Frye SA, et al. The *Mycobacterium tuberculosis* transcriptional landscape under genotoxic stress. BMC Genomics. 2016; 17(1):791.

38. Cortes T, Schubert OT, Banaei-Esfahani A, Collins BC, Aebersold R, Young DB. Delayed effects of transcriptional responses in *Mycobacterium tuberculosis* exposed to nitric oxide suggest other mechanisms involved in survival. Sci Rep. 2017; 7(1):8208.

39. Albrethsen J, Agner J, Piersma SR, et al. Proteomic profiling of *Mycobacterium tuberculosis* identifies nutrient-starvation-responsive toxin-antitoxin systems. Mol Cell Proteomics. 2013; 12(5):1180–1191.

40. Ye J, Coulouris G, Zaretskaya I, Cutcutache I, Rozen S, Madden TL. Primer-BLAST: a tool to design target-specific primers for polymerase chain reaction. BMC Bioinformatics. 2012; 13:134.

41. Somerville W, Thibert L, Schwartzman K, Behr MA. Extraction of *Mycobacterium tuberculosis* DNA: a question of containment. J Clin Microbiol. 2005; 43(6):2996–2997.

42. Votintseva AA, Bradley P, Pankhurst L, et al. Same-day diagnostic and surveillance data for tuberculosis via whole genome sequencing of direct respiratory samples [Internet]. J Clin Microbiol. 2017; 55(5):1285–1298.

43. Gagneux S, DeRiemer K, Van T, et al. Variable host-pathogen compatibility in *Mycobacterium tuberculosis*. Proc Natl Acad Sci U S A. 2006; 103(8):2869– 2873.

44. Freidlin PJ, Nissan I, Luria A, et al. Structure and variation of CRISPR and CRISPR-flanking regions in deleted-direct repeat region *Mycobacterium tuberculosis* complex strains. BMC Genomics. 2017; 18(1):168.

45. Kapopoulou A, Lew JM, Cole ST. The MycoBrowser portal: A comprehensive and manually annotated resource for mycobacterial genomes. Tuberculosis. 2011; 91(1):8–13.

46. Harkins KM, Buikstra JE, Campbell T, et al. Screening ancient tuberculosis with qPCR: challenges and opportunities. Philos Trans R Soc Lond B Biol Sci. 2015; 370(1660):20130622.

47. Li S, Liu B, Peng M, et al. Diagnostic accuracy of Xpert MTB/RIF for tuberculosis detection in different regions with different endemic burden: A systematic review and meta-analysis. PLoS One. Public Library of Science; 2017; 12(7):e0180725.

48. Hegde SR, Rajasingh H, Das C, Mande SS, Mande SC. Understanding communication signals during mycobacterial latency through predicted genome-wide protein interactions and boolean modeling. PLoS One. 2012; 7(3):e33893.

49. Arnvig KB, Comas I, Thomson NR, et al. Sequence-Based Analysis Uncovers an Abundance of Non-Coding RNA in the Total Transcriptome of *Mycobacterium tuberculosis*. PLoS Pathog. Public Library of Science; 2011; 7(11):e1002342.

50. Brown AC, Bryant JM, Einer-Jensen K, et al. Rapid Whole-Genome Sequencing of *Mycobacterium tuberculosis* Isolates Directly from Clinical Samples. J Clin Microbiol. 2015; 53(7):2230–2237.

