## Supplementary Methods for "Towards next generation diagnostics for tuberculosis: identification of novel molecular targets by large-scale comparative genomics"

**Supplementary methods 1. - BLAST searches and databases**

To build the NTM custom database, 4,277 assemblies of NTM species were downloaded and used to build a BLAST database. These assemblies comprised all the *Mycobacterium* genomes deposited in the RefSeq and GenBank databases (June 2017), excluding MTBC species. Both databases were used to search the *M. tuberculosis* genes, using the blastn algorithm (instead of megablast) with a word size (seed) of 7 bp for maximum sensitivity. We found this to be relevant as both local and web-based blast tools use the megablast algorithm with a word size of 28 bp by default, hence only looking for highly similar sequences.

**Supplementary methods 2. - Samples tested in the qPCR experiments**

The assays were carried out using DNA from all MTBC lineages (with exception of lineage 7 due to unavailability), Human DNA, a mock sample with mixed DNA from 20 different bacterial species (ATCC® MSA-1002™ - *A. baumannii, A. odontolyticus*, *B. cereus*, *B. vulgatus, B. adolescentis*, *C. beijerinckii, C. acnes*, *D. radiodurans, E. faecalis*, *E. coli, H. pylori, L. gasseri*, *N. meningitidis*, *P. gingivalis*, *P. aeruginosa, R. sphaeroides, S. aureus*, *S. epidermidis*, *S. agalactiae* and *S. mutans*) and 17 different species of non-tuberculosis mycobacteria: *M. abscessus*, *M. avium*, *M. chelonae*, *M. fortuitum*, *M. gastri*, *M. gordonae*, *M. intracellulare*, *M. kansasii*, *M. lentiflavum*, *M. marinum*, *M. mucogenicum*, *M. peregrinum*, *M. phlei*, *M. scrofulaceum*, *M. smegmatis*, *M. szulgai*, *M. vaccae*.
